# Viral metagenomic analysis of the cheese surface: a comparative study of rapid procedures for extracting virus-like particles

**DOI:** 10.1101/503599

**Authors:** Eric Dugat-Bony, Julien Lossouarn, Marianne De Paepe, Anne-Sophie Sarthou, Yasmina Fedala, Marie-Agnès Petit, Stéphane Chaillou

## Abstract

The structure and functioning of microbial communities from fermented foods, including cheese, have been extensively studied during the past decade. However, there is still a lack of information about both the occurrence and the role of viruses in modulating the function of this type of spatially structured and solid ecosystems. Viral metagenomics was recently applied to a wide variety of environmental samples and standardized procedures for recovering virus-like particles from different type of materials has emerged. In this study, we adapted a procedure originally developed to extract viruses from fecal samples, in order to enable efficient virome analysis of cheese surface. We tested and validated the positive impact of both addition of a filtration step prior to virus concentration and substitution of purification by density gradient ultracentrifugation by a simple chloroform treatment to eliminate membrane vesicles. Viral DNA extracted from the several procedures, as well as a vesicle sample, were sequenced using Illumina paired-end MiSeq technology and the subsequent clusters assembled from the virome were analyzed to assess those belonging to putative phages, plasmid-derived DNA, or even from bacterial chromosomal DNA. The best procedure was then chosen, and used to describe the Epoisses cheese virome. This study provides the basis of future investigations regarding the ecological importance of viruses in cheese microbial ecosystems.

**IMPORTANCE:** Whether bacterial viruses (phages) are necessary or not to maintain food ecosystem function is not clear. They could play a negative role by killing cornerstone species that are necessary for fermentation. But they might also be positive players, by preventing the overgrowth of unwanted species (*e.g*. food spoilers). To assess phages contribution to food ecosystem functioning, it is essential to set up efficient procedures for extracting viral particles in solid food matrix, then selectively sequence their DNA without being contaminated by bacterial DNA, and finally to find strategies to assemble their genome out of metagenomic sequences. This study, using cheese rind surface as a model, describes a comparative analysis of procedures for selectively extracting viral DNA from cheese and to efficiently characterize the genome of dominant phages with cross-sample assembly.

## INTRODUCTION

The cheese surface hosts dense and diverse microbial communities composed of bacteria, yeasts and filamentous fungi. Composition of these communities has been studied for decades (see (1) and (2) for reviews). With the help of high throughput sequencing techniques, we now have detailed pictures of the communities present in a large panel of cheese varieties, and from all over the world (3–6). However, like many other microbial ecosystems, there is still a lack of knowledge on whether and how viral diversity controls the structure of cheese surface microbiota.

Viruses infect all forms of life (7), from prokaryotes (8) to eukaryotes (9) and, in some particular cases, viruses themselves (10). In microbial ecosystems, viral predation is known to greatly influence the structure and functioning of microbial communities (11–13). Nevertheless, since virus genomes lack a single marker sequence for phylogenetic analysis, the structure and role of viral communities in nature are rarely evaluated. Recently, viral shotgun metagenomics helped to describe viral communities from environmental samples including ocean, freshwater, soil, mammalian gut and, to a minor extent, fermented food products (14). Nevertheless, there is currently no literature on the viral diversity of the cheese surface.

Protocols for extraction of virus-like particles (VLPs) have been optimized for different environments (15–18). For food samples, most available procedures have been designed for the recovery of foodborne viruses potentially affecting human health such as noroviruses, rotaviruses (RoV) and hepatitis viruses (19), but not for microbial viruses such as bacteriophages. Furthermore, because the cheese surface has peculiar characteristics such as the presence of caseins at high concentration, high fat and salt content, it proves necessary to set up a dedicated procedure. Ideally, the method should be easy-to-use and rapid enough to be compatible with medium (dozens to hundred samples) to large-scale studies (hundreds to thousands samples) such as those performed to describe microbial communities (6).

Taking into account all these constraints, we compared four procedures for the isolation of viruses from cheese surfaces. The backbone of the protocol followed the PEG-based protocol already evaluated by Castro-Mejia et al. on fecal samples (15). However, we replaced the buffer used for sample suspension, and tested both the effect of adding a filtration step before PEG precipitation in order to completely remove microbial cells and the substitution of the expensive and time-consuming density gradients for VLPs purification by a simple chloroform treatment which is used for membrane vesicles removal (20, 21) (Fig. 1). We evaluated the usefulness of the four procedures for cheese virome analysis (limited to the viruses with DNA genomes, expectedly the most abundant) using three different types of cheese, namely Camembert, Epoisses and Saint-Nectaire, and following two criteria: particles recovery and particles purity. Finally, we assessed the level of microbial DNA contamination in virome sequencing data for one type of cheese.

**FIG 1.**
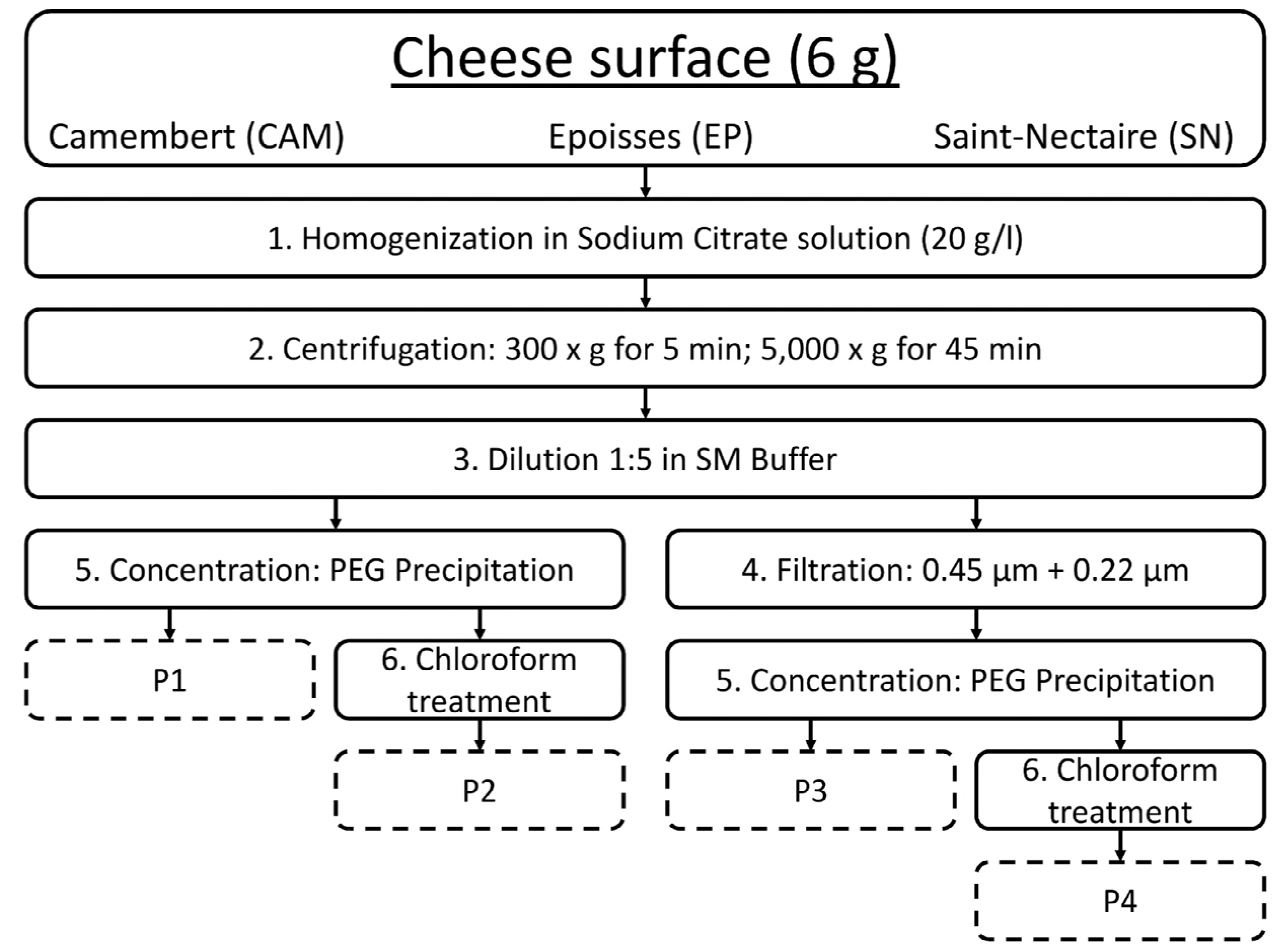
Schematic representation of the experimental procedure for the extraction of virus-like particles from cheese. The surface of three types of cheese was processed according to four routes named P1 to P4. Steps 1 (homogenization), 2 (centrifugation), 3 (dilution) and 5 (concentration) are common to all protocols, contrary to steps 4 (filtration) and 6 (chloroform treatment) which are optional. For each cheese type, three biological replicates (independent cheeses purchased at the same date and from a unique producer) were performed leading thus to 12 samples per cheese-type.

## RESULTS

### Quantification and purity of VLPs recovered from the cheese surface

Among the four tested procedures, only those containing chloroform treatments (P2 and P4) resulted in VLP solutions sufficiently pure to allow nanoparticle counting using the interferometric light microscope (Table 1). We observed 1 × 10^9^ to 4 × 10^10^ nanoparticles per gram of cheese surface, and filtration did not induce a drastic loss of particles. Counts were higher in Epoisses and Camembert than in Saint-Nectaire cheese. Camembert had the highest nanoparticles per microbial cell ratios. This type of cheese had the lowest bacterial counts (Table S1), whether such a community leads to higher cell lysis mediated by viruses or higher levels of membrane vesicles remains to be investigated. VLP solutions prepared using procedures P1 and P3, without chloroform treatment, were very dense, milk-white and led to hard noise in interferometer's films with a myriad of spots larger than those typically observed for viruses. Microbial cells clearly contaminated the VLP fractions when using procedures P1 and P2, without filtration (Table 1, column 3). It has to be mentioned we also observed bacterial cells in one filtrated sample, *i.e*. Camembert cheese with P3, indicating probable post-filtration contamination.

**Table 1.** Quantification of VLPs using interferometry and microbial cells using plate numbering.

| Type of Cheese and of protocol | Total nanoparticles (log per g) (SD) | Presence of cells | Presence of noise | Total biomass (log CFU per g) (SD) | Ratio nanoparticles / cell |
| --- | --- | --- | --- | --- | --- |
| **CAM P1** | NA | +++ | +++ | 8.71 (0.11) | NA |
| **EP P1** | NA | +++ | +++ | 10.10 (0.31) | NA |
| **SN P1** | NA | +++ | +++ | 10.42 (0.07) | NA |
| **CAM P2** | 10.38 (0.01) | ++ | - | 8.71 (0.11) | 47.29 (11.91) |
| **EP P2** | 10.18 (0.23) | + | - | 10.10 (0.31) | 1.88 (2.07) |
| **SN P2** | 10.04 (0.24) | + | - | 10.42 (0.07) | 0.66 (0.14) |
| **CAM P3** | NA | ++ | +++ | 8.71 (0.11) | NA |
| **EP P3** | NA | - | +++ | 10.10 (0.31) | NA |
| **SN P3** | NA | - | +++ | 10.42 (0.07) | NA |
| **CAM P4** | 10.00 (0.13) | - | - | 8.71 (0.11) | 21.77 (12.15) |
| **EP P4** | 10.13 (0.37) | - | - | 10.10 (0.31) | 2.40 (3.35) |
| **SN P4** | 9.59 (0.40) | - | - | 10.42 (0.07) | 0.17 (0.10) |

NA: not available because of noise in interferometric measures

Two-layer iodixanol gradients were used as quality controls in order to separate microbial cells, debris and membrane vesicles from viruses, enabling to visually observe the effect of the four procedures on the quantity and purity of VLP solutions (Fig. 2). For the three types of cheese tested, similar profiles were observed after ultracentrifugation. A strong band located at the top of the lightest density layer was present in samples prepared without filtration and chloroform treatment (P1) suggesting high contamination with microbial cells, debris and membrane vesicles (Fig. 2, red arrow). This band was still present – albeit slightly less intense – in filtered samples (P3), suggesting that cheese rind is rich in membrane vesicles. Finally, this band was completely absent from samples prepared with procedures including chloroform treatment (P2 and P4) confirming the efficiency of such treatment for removing membrane vesicles from VLP solutions (20). A visible band containing viruses formed at the proximity of the 45% iodixanol cushion (Fig. 2, blue arrow), at a density corresponding to 40% iodixanol, as estimated with a refractometer. This virus band was barely visible, however, in samples treated with chloroform (P2 and P4) when compared to untreated samples (P1 and P3). Nanoparticles quantification using interferometry in the viral band of Epoisses samples after dialysis indicated that samples treated with cholorofom (P2 and P4) contained approximately ten times less nanoparticles than untreated samples (P1 and P3). For Camembert cheese and procedure P3, we observed two distinct bands near the 45% iodixanol cushion. Transmission electronic microscopy revealed that viruses were more abundant in the lowest one.

**FIG 2.**
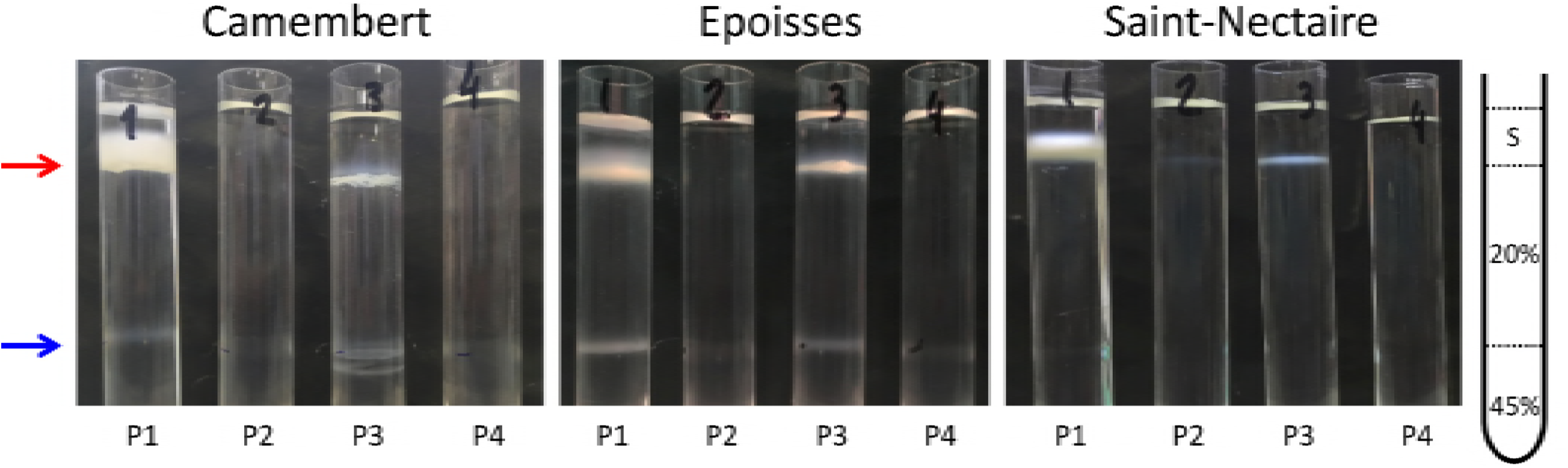
Photographs of iodixanol density gradients after ultracentrifugation of cheese VLP solutions. Cellular debris and membrane vesicles are concentrated at the top of the 20% iodixanol layer (red arrow) and VLPs at the top of the 45% iodixanol layer (blue arrow). A schematic representation of the gradient is shown on the right. S: sample.

Altogether, these results suggest that filtration is an essential step in cheese VLPs preparation in order to completely eliminate microbial cells, which represent the major source of DNA contamination in virome studies. Furthermore, chloroform treatment is very beneficial for VLP solutions’ cleaning and removal of membrane vesicles, which might contain pieces of microbial DNA (see below), but at the cost of reducing the nanoparticle recovery.

### Effect of the extraction procedure on the composition of the Epoisses virome

Saint-Nectaire exhibited the lowest nanoparticles per cell ratio and Camembert the highest, Epoisses being an intermediate situation (Table 1). Furthermore, plate counting indicated a higher bacterial density in Epoisses cheese than in Camembert (Table S1), bacteria being expected as the main targets of microbial viruses (bacteriophages or phages) in the cheese ecosystem. For those two reasons, we selected Epoisses cheese as the best candidate for further virome analysis. DNA was successfully extracted from VLP solutions produced from Epoisses cheese using the four procedures and prior the gradient step. DNA yields are available in Table S2. DNA samples were then sequenced, in order to assess the impact of both filtration and chloroform treatment on the final sequence dataset (see Material and Methods section). In addition, for sample P3, the vesicle band was recovered from iodixanol gradient, DNA was extracted using the same protocol as viruses, and sequenced. The virome sequence data from each of the 12 samples (4 protocols x 3 biological replicates) and that of the vesicles sample were binned together for assembly of the virome (see Material and Methods section for more details).

Quality filtered reads were pooled and assembled into 910 contigs of length greater than 2,500 bp, a threshold used for keeping putative small virus genomes (22). Of these, 124 contigs were selected for further analysis based on VirSorter results and coverage criteria (see Material and Methods section). These 124 contigs were ranging in size from 2.5 kb to 122 kb with a N50 of 24.4 kb and were designated as the Epoisses cheese virome. Mapping of the quality filtered reads against this virome revealed that microbial cell contamination was higher with protocol P1 and in the vesicles sample than in any of the other three protocols (Fig. 3).

**FIG 3.**
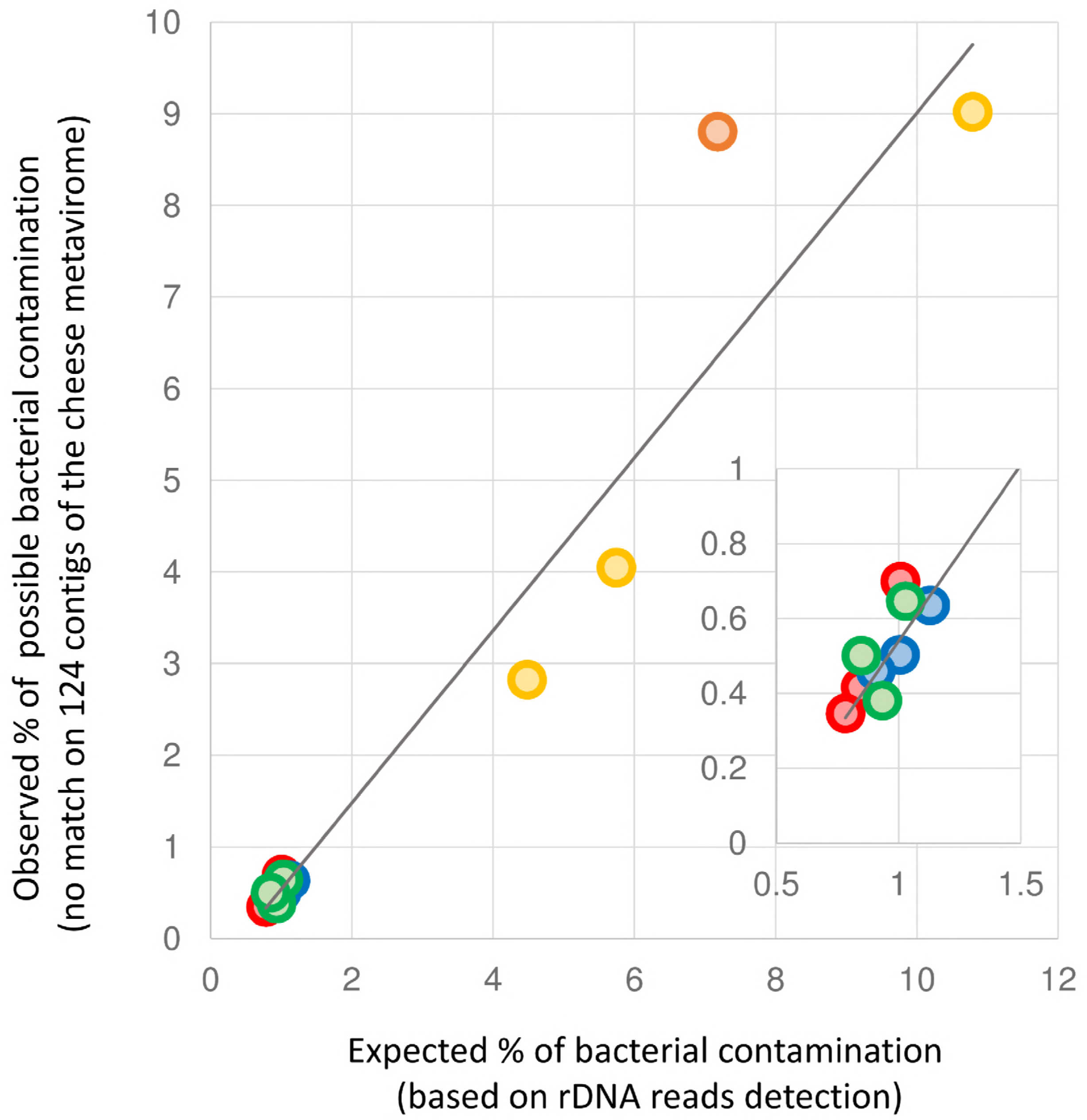
Comparative plot showing the estimated level of bacterial DNA contamination in the Epoisses cheese virome. The expected percentage (x axis) is based on the % of rDNA reads detected by SortMeRNA software. This value was used to calculate the estimated % of bacterial chromosomal DNA using a multiplying factor of (x100) based on estimation that several copies of rRNA operons (*e.g*. 5 copies of ~5 kb in average) would represent about 1% of a bacterial chromosome of an average size of 2.5 Mb. The observed % of possible bacterial contamination was obtained by looking at the % of reads not matching the 124 contigs of the cheese virome. Yellow (P1), red (P2), green (P3), blue (P4) and orange (vesicles sample).

The contigs were further analyzed and separated into four classes based on PHASTER and Blast results (*i.e*. probable phage contigs, plasmid-derived contigs, putative plasmid-derived contigs and unclassified contigs) (Dataset S1). Principal coordinate analysis performed on the Bray-Curtis dissimilarity index calculated from the abundance table of the 124 contigs (Fig. 4A) showed an ordination of the samples according to the VLP extraction procedure, with samples obtained with P1 and P3 being separated from samples obtained with P2 and P4. This indicated a possible impact of the chloroform treatment on the quality of the resulting virome. Datasets produced with protocols P2, P3 and P4 were strikingly enriched for reads mapping to probable phage contigs (>99% in average) when compared to VLP fraction obtained with P1 (94% in average) (Fig. 4B), which was largely explained by the higher proportion of unmapped reads present in P1 samples (Fig. 4F). The relative proportion of reads mapping to the 19 plasmid-derived contigs was greatly reduced in samples treated with P3 (filtration) and even more so in samples treated with P2 (chloroform) and P4 (chloroform and filtration), when compared to P1 (Fig. 4C), reflecting the positive impact of both filtration and chloroform treatment on the quality of the final virome. The relative proportions of reads matching putative plasmid-derived contigs (Fig. 4D) and unclassified contigs (Fig. 4E) remained similar regardless of the VLP extraction procedure.

**FIG 4.**
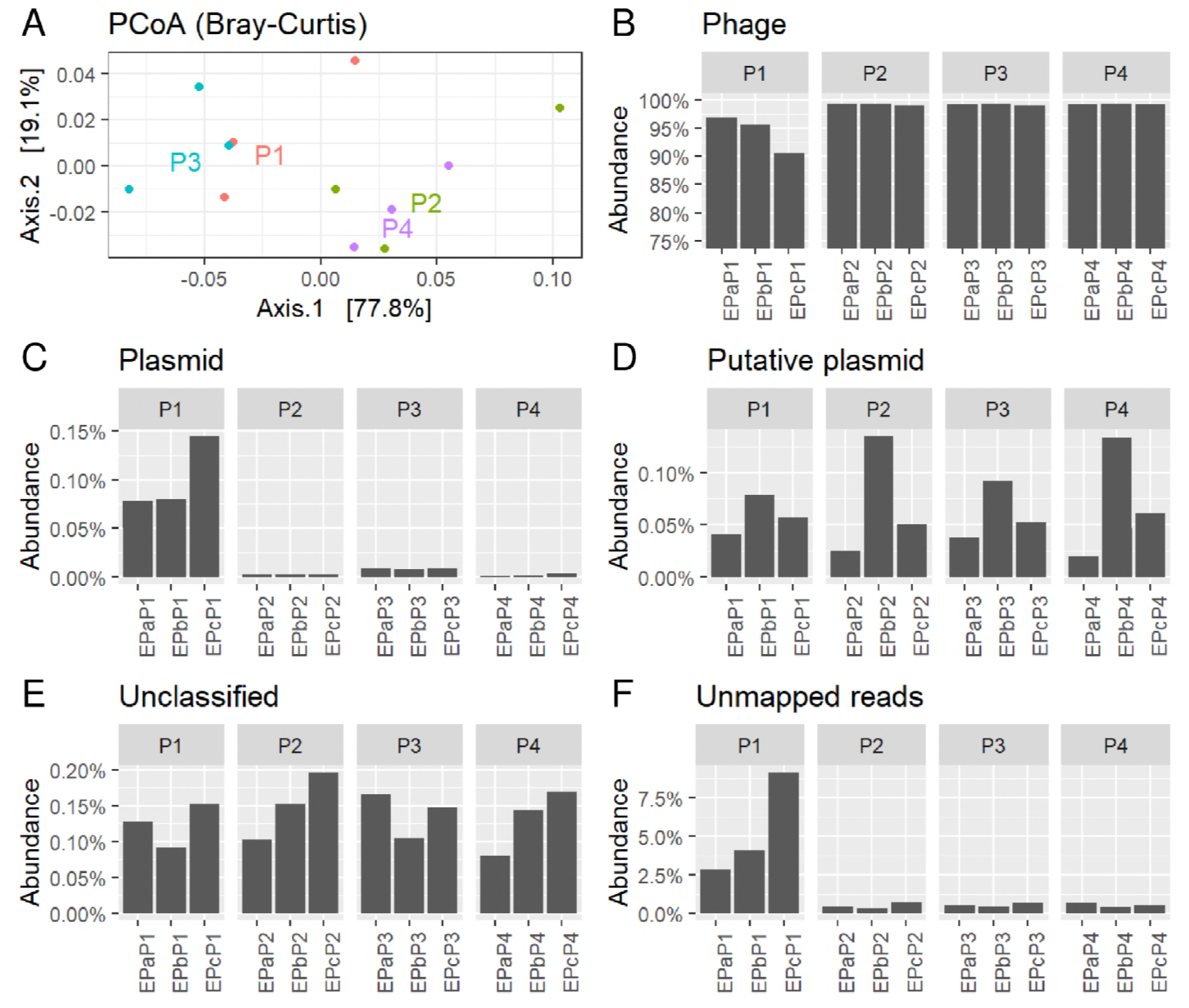
Differences in the Epoisses cheese virome composition according to the VLP extraction procedure. Principal coordinate analysis based on the Bray-Curtis dissimilarity index (A). Relative abundance of the contigs annotated as probable phages (B, 72 contigs), plasmids (C, 19 contigs), putative plasmids (D, 20 contigs), unclassified (E, 14 contigs) and unmapped reads (F).

Detailing the composition of each virome evidenced that the detection of contigs identified as probable phages was not affected by the VLP extraction procedure (Fig. 5), suggesting that neither the filtration step nor the chloroform treatment biased the sample composition. The major differences were observed for the 39 contigs belonging to the categories plasmid and putative plasmid. Most of them were undetected or sporadically detected in samples treated with chloroform (P2, P4). They were on the contrary much higher with protocols P1 (and to a lesser extend P3) and highly abundant in the vesicles sample (39% of the reads). Interestingly, NODE-134, the most abundant contig detected in the vesicles sample (24 *%* relative abundance) and assigned to the plasmid-derived contig category, was only marginally detected in samples treated with protocols P1, P2, P3 and P4 (<0.005% relative abundance).

**FIG 5.**
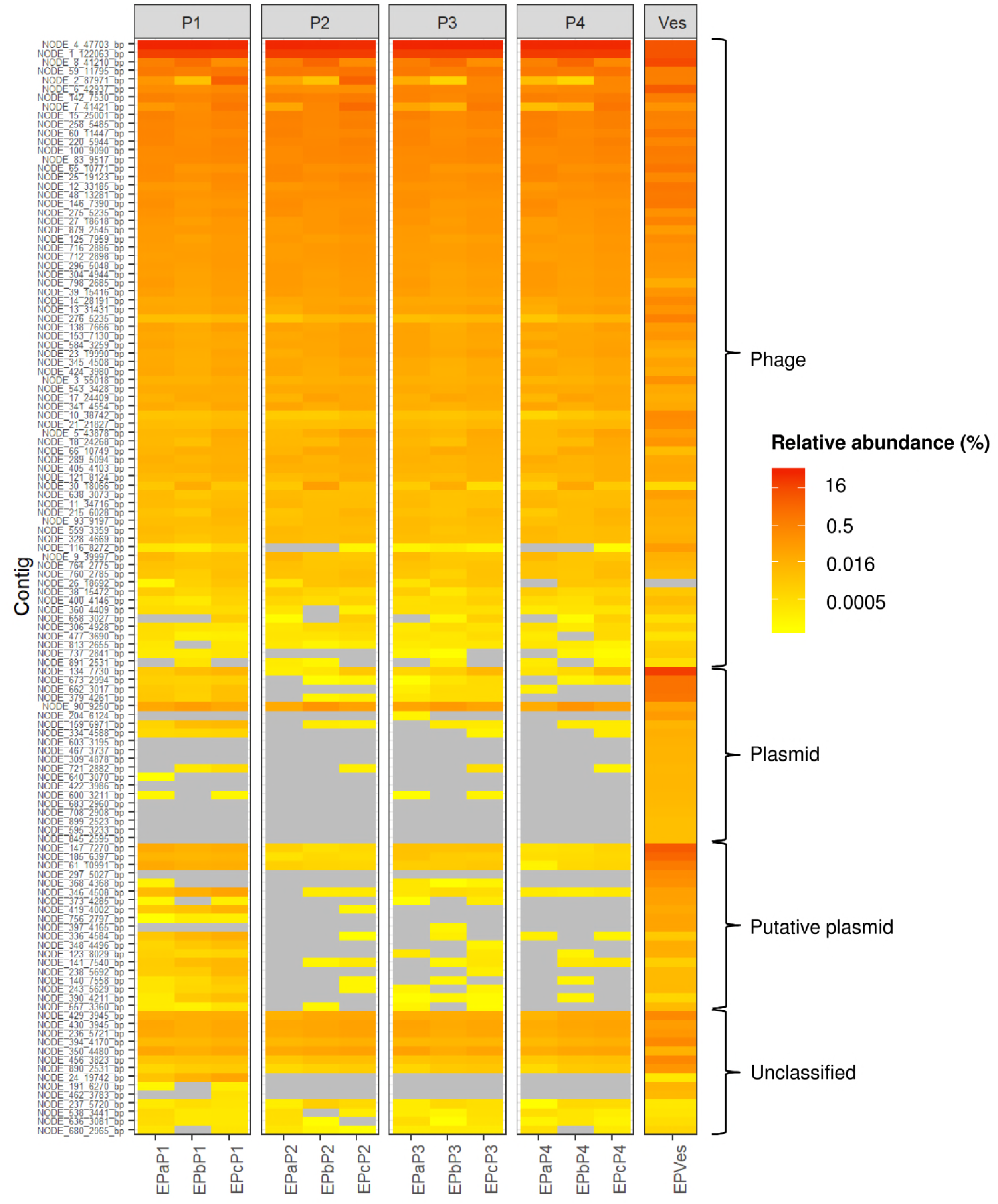
Heatmap of the 124 contigs composing the Epoisses virome. Samples were grouped according to the VLP extraction procedure. Contigs were separated into four categories (phage, plasmid, putative plasmid and unclassified) according to the annotation results and sorted by abundance inside each group. Yellow to red: lowest to highest relative abundance. Grey: not detected. Ves: vesicles sample.

### Composition of the Epoisses cheese virome

Three repeats of the extraction-sequencing on Epoisses cheese bought simultaneously from the same manufacturer gave highly reproducible results. Two contigs, NODE-1 and NODE-4, now named Epvir4 and 949_Epvir1, were prevalent, representing relative mean abundances of 63% and 28%, respectively. Epvir4 genome is 42 kb long. It was assigned by VirSorter as a category 2 phage (« quite sure ») but did not share similarity with already known phages. The annotation by RAST, HHpred and BLAST revealed an organization into functional modules (Fig. S1A). Amongst the ORFs with predicted homologues in nr database, most best BlastP matches (63%) were found in the genus *Glutamicibacter* (formerly *Arthrobacter*) suggesting that the host of this abundant phage might be a *Glutamicibacter* species (Dataset S2). 949_Epvir1 genome (Fig. S1B) is 122 kb long and shares 95% nucleotide identity and 95% coverage with *Lactococcus lactis* phage 949, a virulent phage (despite two predicted integrase genes) isolated from cheese whey in New Zealand (23). The third most abundant contig (1.1% of the reads), Epvir8, has a genome 41 kb in length, and shares a small 1.1 kb region with 88% nucleotide identity with *Pseudoalteromonas* phage SL25 (24). Amongst the ORFs with predicted homologues, more than half (56%) were found in *Pseudoalteromonas* phages while another quarter (24%) were found in *Vibrio* phages (Dataset S2 and Fig. S1C)). All other contigs exhibited a relative abundance lower than 1%, most of them being difficult to assign to a putative host. Nevertheless, few contigs were closely related to *Lactococcus lactis* phages (*e.g*. NODE-15, NODE-59, NODE-142), *Pseudoalteromonas* phages (*e.g*. NODE-48, NODE-276), *Vibrio* phages (*e.g*. NODE-12, NODE-116), *Leuconostoc* phages (*e.g*. NODE-18, NODE-658) and *Halomonas* phage (NODE-636). All of these bacterial genera are frequently detected as part of cheese surface (3). The best assignments carried out for all contigs are available in supplementary Dataset S1.

### DNA content of Epoisses membrane vesicles

Much less DNA was present in membrane vesicles, compared to total viromes (Table S2). This has to be kept in mind when comparing the vesicle line to the virome lines in Fig. 5, since colors reflect only relative abundances. Almost 51% of the reads from the vesicles sample mapped to contigs assigned to potential phages, which indicated that phage DNA was present in the vesicles. Interestingly, the reads corresponding to plasmid-derived contigs were specifically enriched in membrane vesicles (10% of total), relative to the viromes (0.03%). It seems therefore that the plasmid content of samples P1 and P3, observed in Fig. 5, originate from membrane vesicles, which are still present since no chloroform treatment was applied in these protocols. The total DNA content obtained from the vesicles fraction was low, relative to viromes, with only 3.8 ng (Table S2), and corresponded to an estimated amount of 1.3 × 10^11^ nanoparticles according to interferometry measures. Supposing all DNA content is plasmidic, with an average size of 5 kb for the plasmids (and a corresponding weight of 0.55 × 10^−17^ g), 3.8 ng of DNA would amount to 7 × 10^8^ plasmidic DNA molecules, suggesting that only 0.5% of the vesicules effectively contain a DNA molecule. The same calculation for phage DNA (~50 kb) would result in 10-fold less DNA containing membrane vesicles.

## DISCUSSION

With the increasing attraction for virome sequencing, protocols for extracting viruses from diverse environmental samples have been developed during the past decade. Prior to nucleic acid extraction and sequencing, these protocols usually included sample pretreatment to make viral particles accessible for extraction, virus concentration and finally virus purification (18). Each step has to be adapted, according to the type of samples studied and the type of virus targeted. For cheese, we chose to blend the samples after a dilution step in trisodium citrate, in order to maximize the chance for recovering viral particles from the matrix. Indeed, citrate is a complexing agent for calcium and allows casein solubilization. It is largely used in nucleic acid extraction protocols for recovering microbial cells from casein network in milk and cheese (25, 26).

We also decided to evaluate the potential benefit of adding a filtration step (0.22 μm) on the quality of VLPs preparation and of the resulting virome. Filtration is generally avoided in virus extraction protocols from environmental sources because some viruses can be very large, even larger than microbial cells (27). Nevertheless, our choice was motivated by the fact that (i) such large viruses are generally hosted by organisms which are not part of the cheese ecosystem such as amoebas, protists and microalgae (ii) it ensures complete removal of microbial cells which generally account for the major source of contaminating DNA sequences in virome studies (28). Our results demonstrated that adding filtration to the extraction procedure reduced microbial contamination of the cheese VLP solutions by microbial cells, without major modification neither in the particle counts nor in the final virome profile.

The best way to purify VLP includes a density gradient ultracentrifugation step (18), in which viruses are separated from other components of the extract based on their physical properties. This technique is, however, very time-consuming, expensive and requires specific technical skills and lab equipments. Chloroform treatment represents a possible alternative for rapid and efficient virus purification (20, 29) and has already been used in the viral metagenomic context (30). This solvent permeabilizes the membranes of bacterial cells and membrane vesicles, disrupting their structural integrity and making nucleic acids available for digestion by nucleases. Our results indicate that such treatment, in combination with filtration, is mandatory for successful quantification of VLP extracted from cheese samples by interferometry. Even if viral capsids are said to be resistant to chloroform (20, 31), it is important to mention that some viruses, in particular enveloped viruses, might be lost after such treatment (18, 20, 21, 32). Comparing the Epoisses cheese VLP solutions obtained with procedures including chloroform treatment or not, indicated a loss of VLP upon chloroform treatment (as illustrated in Fig. 2). Analyzing the nanoparticle counts in the viral band of one Epoisses cheese after density gradient ultracentrifugation and dialysis indicated that we recovered approximately ten times less nanoparticles in chloroform-treated samples versus non-treated samples. However, this was accompanied by a limited reduction of the final DNA yields (<3 times less DNA in average for treated versus untreated samples, Table S2) and we did not observe qualitative differences in the abundances of viral contigs as a function of chloroform treatment. Furthermore, adding chloroform treatment to the extraction procedure resulted in a significant enrichment of the virome in actual viral sequences while drastically reducing the proportion of plasmids and sequences from microbial origin (unmapped reads). For one Epoisses, sequencing of the DNA content of a membrane vesicle fraction, obtained after separation from phage particles on an analytical density gradient, was performed. A low amount of DNA was recovered, in comparison with total viromes, suggesting that at most a few percent of the membrane vesicles contain DNA. This is in line with the current view that the transport of nucleic acids only constitutes one of many potential roles for membrane vesicles. Membrane vesicles are involved in exchanges between cells in the tree domains of life (33). In the bacterial domain, they have multifaceted roles including the delivery of autolysins, cytotoxins and virulence factors, as well as nucleic acids (34, 35). In our membrane vesicle sample, DNA sequencing leads us to formulate several suppositions: i) since half of vesicle reads map to contigs classified as potential phages, it may be that Epoisses membrane vesicles serve as decoys and trap phages, as recently reported in *Vibrio cholerae* (36). The interaction between phage tail fibers and membrane vesicles harboring proper phage receptors could contribute to the emptying, and thereby inactivating of phage virions, after DNA injection. ii) The enrichment in reads corresponding to predicted plasmids, compared to total viromes, suggests that Epoisses membrane vesicles transport plasmid DNA, as described for an *Escherichia coli* strain (37). iii) A third category of DNA content in this vesicle fraction, composed of circular contigs that might be plasmidic but with no hit in databases, might not belong to vesicles. Indeed, the relative proportions of these contigs are similar in each virome, irrespective of the chloroform treatment. These DNA fragments might be packaged by transducing phage particles, which are more resistant to chloroform than vesicles. One has to suppose that such transducing particles have a lower density than *bona fide* phage particles to explain their presence together with vesicles fraction.

The set up of the protocol most adapted to recover viral communities from cheese rind was considerably facilitated by the use of a microscopic device called interferometer, allowing evaluating in real time the total nanoparticle counts (38, 39). It permitted in particular to choose, among several cheeses, those with higher nanoparticle titers for the present study. Camembert and Epoisses cheeses had markedly more nanoparticles, compared to Saint-Nectaire (Table 1). Camembert and Epoisses cheese are both soft cheeses, characterized by a bloomy and washed rind, respectively. These types of cheese typically have a higher moisture content than semi-soft pressed cheese with natural rind such as Saint-Nectaire (40, 41). Although a more comprehensive comparison including more cheese varieties would be required, our results are congruent with the general assumption that viruses would be more abundant in hydrated environment, compared to dry ones.

The rapid procedure presented in this study enabled to obtain sufficient quantity of viral DNA from Epoisses cheese (equivalent of 1.5 gram as starting material for each protocol) for direct library preparation prior to sequencing, avoiding the use of multiple displacement amplification (MDA) (DNA quantities obtained for all samples presented in this study are available in Table S2). MDA is widely used in viral metagenomic studies due to the insufficient DNA material extracted from the samples (18). However, amplification bias has been documented for this technique (42) which might provoke distortion in the viral community profiles and making preferable direct DNA sequencing when possible.

The Epoisses virome analyzed in this study was composed of many contigs sharing high sequence identity with known *Lactococcus lactis* phages including the widely known 936, P335 and C2-like groups. Interestingly, the second most abundant one (28% of reads), named 949_Epvir1, was very similar to the *L. lactis* phage 949, a virulent phage isolated from cheese whey in New Zealand, which is phylogenetically distant from those groups of commonly isolated *L. lactis* phages (23). *Lactococcus lactis* is used as starter in the manufacture of Epoisses cheese. In a previous work describing the microbial diversity in twelve french cheese varieties, it was detected as the dominant bacterial taxa in the core of Epoisses cheese and was also highly detected in the rind (3). In the same study, other dominant bacterial genera of the Epoisses cheese rind included not-deliberately inoculated taxa such as, by order of importance, *Psychrobacter, Marinomonas, Vibrio, Pseudoalteromonas, Glutamicibacter, Mesonia, Enterococcus, Lactobacillus* and *Halomonas*. Interestingly, phages potentially infecting some of such non-starter bacterial taxa, including *Glutamicibacter* (Epvir4, most abundant contig in the Epoisses virome, 63% of the reads), *Pseudoalteromonas* (Epvir8, third most abundant contig, 1.1% of the reads), *Vibrio, Leuconostoc* and *Halomonas*, were also detected in the Epoisses cheese virome.

The cheese surface microbiota is in constant evolution during the ripening process, and is characterized by the successive development of different microbial groups. Microbial interactions between different species have been observed in cheese (43–46), providing the first elements of comprehension regarding biotic forces sustaining microbial assemblage in this peculiar environment. Our results indicate that many microbial species living on the cheese surface are also subjected to viral predation, and shed light on the need of careful evaluation of the impact of viruses on the dynamic of the cheese microbial ecosystem.

## CONCLUSION

In this study, we optimized a rapid protocol for VLP extraction from the cheese matrix suitable for subsequent virome analysis. We demonstrated its efficiency by extracting VLPs from three different type of cheese and produced the first cheese surface virome using Epoisses cheese as a model. Our results emphasized the positive impact of both filtration and chloroform treatment on the final virome quality in the cheese context. In the future, we anticipate virome analysis will expand our knowledge on cheese microbial ecosystems and possibly give the opportunity to better understand the rules of microbial assembly that occur in this fermented food.

## MATERIAL AND METHODS

### Sampling procedure

Three types of French surface-ripened cheese were studied, namely Camembert (CAM, bloomy rind), Epoisses (EP, washed rind) and Saint-Nectaire (SN, natural rind). For each type of cheese, three cheeses produced at the same date and from a unique producer were purchased and analyzed as replicates. Rind was gently separated from the core using sterile knives, and mixed using a blender.

### Microbiological analysis

One gram of the cheese surface was diluted 1:10 in sterile saline solution (9 g/l NaCl) and homogenized with an Ultra Turrax Homogenizer (Labortechnik, Staufen, Germany) at full speed for 1 min. Serial dilutions were performed in 9 g/l NaCl and microorganisms were enumerated by surface plating in duplicate on specific agar base medium. Cheese-surface bacteria were enumerated on brain heart infusion agar (Biokar Diagnostics) supplemented with 22.5 mg/l amphotericin B after 3 to 5 days of incubation at 25°C under aerobic conditions. Lactic acid bacteria were enumerated on de Man-Rogosa-Sharpeagar (pH 6.5, Biokar Diagnostics) supplemented with 22.5 mg/l amphotericin B after 3 days of incubation at 30°C under anaerobic conditions. Finally, fungal populations were enumerated on yeast extract-glucose-chloramphenicol (Biokar Diagnostics) supplemented with 2,3,5-triphenyltetrazolium chloride (10 mg/l) after 3 to 5 days of incubation at 25°C under aerobic conditions.

### Extraction of virus-like particles (VLPs)

Four protocols, named P1, P2, P3 and P4, were tested in parallel starting from the same material (Fig. 1). Six grams of cheese rind was diluted 1:10 with cold trisodium citrate (2% w/v) into a sterile bag and mixed for 1 min using a BagMixer (Interscience). Citrate is a complexing agent for calcium and allows casein solubilization. It is largely used in nucleic acid extraction protocols for recovering microbial cells from casein network in milk and cheese (25, 26). The solution was centrifuged at 300 × g for 10 min at 4°C in order to pellet big aggregates. The supernatant was centrifuged at 5,000 × g for 45 min at 4°C in order to pellet microbial cells. At this stage, the supernatant containing free VLPs was diluted 1:5 with cold SM buffer (200 mM NaCl, 10 mM MgSO_4_, 50 mM Tris pH 7.5) and split, half being used for filtration-free protocols (protocols P1 and P2) and half being successively filtrated using 0.45μm and 0.2μm polyethersulfone membranes and glass vacuum filter holders (Millipore) (protocols P3 and P4). Samples, filtrated or not, were then supplemented with 10% PEG 6,000 (Sigma) and kept at 4°C over-night after dissolution for VLPs precipitation. After centrifugation at 6,000 × g for 1h at 4°C, pellets were resuspended with 2 ml of cold SM buffer and split again, half being stocked at 4°C (protocols P1 and P3) and half being treated with chloroform in order to eliminate membrane vesicles (protocols P2 and P4). More specifically, 1 volume of fresh, non-oxidized chloroform was added to the sample and mixed thoroughly for 1 min using a vortex to create an emulsion. After centrifugation at 15,000 × g for 5 min at 4°C, the aqueous phase containing viral particles was recovered.

### Particles counting using Interferometry

A “home-made” interferometric light microscope (ILM) (38) was used to count nanoparticles (i.e. both vesicles and viruses) present in our VLP fractions as previously described (39). Briefly, 5 μl of each sample were used to collect a stack of 200 images (CMOS camera). We developed a simple ImageJ (47) script that allows first background subtraction and image quality enhancement, then particles localization on each image. An average number of nanoparticles per frame was obtained. To calibrate the concentration estimates, crude lysates of phages P1, T4, T5, T7, λCI857 and φX174 were titrated on *Escherichia coli* MG1655, and counted with the ILM device. These counts were converted into concentrations, assuming that all particles present in the observation volume (10^−8^ ml) were detected with the ILM device. Table S3 and Fig. S2 showed a good match between the two measurements, usually within a +/− 3 fold range for phages P1, T4, T5, T7 and λCI857, whereas φX174 was underestimated by 20 fold. We next compared the ILM estimates with epifluorescence microscopy on two VLP samples obtained from Epoisses cheese (Fig. S3). As shown in the result section, virome samples of Epoisses rind also contain vesicles, which are detected by ILM, but not by epifluorescence, since they mostly do not contain DNA. In accordance with expectations, 1.5 to 1.9-fold more particles were counted by ILM than by epifluorescence. We concluded that the ILM measurements give reasonable estimates of nanoparticle concentrations (both vesicles and VLP) of viromes.

### Particles purity evaluation using iodixanol gradients

Two-layer iodixanol discontinuous gradients of 45% and 20% (w/v) in SM Buffer (50 mM Tris pH7.5, 100 mM NaCl, 10 mM MgCl2) were prepared by adding first 6 ml of the 20% fraction in a 12 ml ultracentifugation tube, and then under layering 4.5 ml of the 45% fraction in the bottom of the tube, using a glass transfer pipette and a pipette pump. Each VLP solution (1 ml diluted 1:2 with SM Buffer) was finally added on top of the gradient and the tubes were centrifuged at 200,000 × g for 5 h at 10°C, in an SW41 rotor, using a Beckman XL-90 ultracentrifuge.

### Viral DNA extraction

500 μl of the VLPs preparation were treated for 30 min at 37°C with 1 U of TURBO DNase (Invitrogen) in order to digest free DNA. DNase was inactivated by the addition of 5.5 μl of 100 mM EDTA and the sample was placed on ice for 5 min. The entire content of the tube was transferred to a 2 ml tube containing a gel that improves separation between the aqueous and organic phases (Phase Lock Gel Heavy; Eppendorf AG, Hamburg, Germany). One volume of phenol-chloroform (1:1; saturated with 10 mM Tris pH 8.0, and 1 mM EDTA) was added to the sample and, after gentle mixing for 1 min by inversion, the tube was centrifuged at 10,000 × g for 10 min at 4°C. The aqueous phase was then transferred to another Phase Lock Gel tube. After adding one volume of phenol-chloroform and gentle mixing for 1 min by inversion, centrifugation was performed at 10,000 × g for 10 min at 4°C. The aqueous phase was then transferred to another Phase Lock Gel tube. After adding one volume of chloroform and gentle mixing for 1 min by inversion, centrifugation was performed at 10,000 × g for 10 min at 4°C. The aqueous phase (approximately 300 μl) was recovered in a 1.5 ml tube and DNA was precipitated by adding two volumes of absolute ethanol, 50 μl of sodium acetate (3 M, pH 5.2) and 3 μl of glycogen (5 mg/ml; Invitrogen) as carrier. After incubation on ice for 10 min, the DNA was recovered by centrifugation at 13,000 × g for 30 min at 4°C. The DNA pellet was subsequently washed with 800 μl of 70% (vol/vol) ethanol. After centrifugation at 13,000 x g for 10 min at 4°C, the DNA pellet was dried at room temperature for 30 min and dissolved in 20 μl of 10 mM Tris pH 7.5. Purified DNA was quantified using the Qubit DNA Broad Range assay (ThermoFisher Scientific).

### Viral DNA sequencing and bioinformatic analysis

Library preparation and sequencing were performed at the GeT-PlaGe platform (Toulouse, France). Briefly, libraries were prepared using the NEBNext^®^ Ultra™ II DNA Library Prep Kit (New England Biolabs) and sequencing was performed on the MiSeq plateform (Illumina) according to the manufacturer’s instructions. Raw reads were quality filtered using Trimmomatic (Bolger et al., 2014) in order to remove paired reads with bad qualities (with length < 125 bp, with remnant of Illumina adapters and with average quality values below Q20 on a sliding window of 4 nt). Furthermore, only the remaining paired reads were kept while those being unpaired at the end of the filtering process were kept aside. The overall filtering process yielded an average of 1.36 million paired-end reads (2 × 250 bp) per sample (2.72 million reads total). All quality-filtered reads from the various experiments were pooled in order to proceed to only one global assembly. Therefore, reads from similar phages but present in several samples would be binned together during the assembly process. Assembly was performed using SPAdes with the *meta* option and increasing *kmer* values -k 21,33,55,77,99,127 (48). Contigs with length below 2,500 bp, unlikely to encode complete viruses, were discarded. The resulting 910 filtered contigs were first analyzed with VirSorter (49), which returned 92 putative viral contigs. A list of “most abundant and pertinent” contigs was constructed using three criteria: 1) detected as viral with Virsorter, after filtering out those placed in categories 3 or 6 (not so sure) with coverage below 10 (remaining 78 contigs), 2) detected as circular contig, even if not detected as viral (adds 34 contigs) 3) coverage above 100, even if not detected as viral (final list of 124 contigs). These contigs were further analyzed using PHASTER (50) and compared to both the Viruses section of nucleotide database at NCBI and to the complete database using Blast (51) in order to provide a first characterization of putative phage-encoding contigs. Then, quality filtered reads (both paired and unpaired) were mapped individually for each sample against the potential viral contigs using Bowtie2 aligner with default values (52) in order to produce an abundance table. This table was processed with the R package Phyloseq for statistical analysis and data visualization (53). The level of bacterial DNA contamination in the cheese viromes was estimated by detection of ribosomal DNAs among the reads using SortMeRNa software (54) and the SILVA v129 database (55). The three most prevalent contigs present in the final virome, namely Epvir4, 949_Epvir1 and Epvir8, were further annotated using RAST (56). Homologs of the predicted proteins in the contigs were additionally compared to the NCBI nr database using BlastP and HHpred (57) with an E-value cutoff of 10^−8^.

### Accession numbers

Raw sequence data were deposited at the Sequence Read Archive of the National Center for Biotechnology Information under the accession numbers SRR8080803 to SRR8080815 (bioproject PRJNA497596).

## ACKNOWLEDGMENTS

This work was conducted in the frame of the Virome Access project supported by the French National Institute for Agricultural Research (INRA) and the Meta-omics and Microbial Ecosystems (MEM) programme. The funders had no role in study design, data collection and interpretation, or the decision to submit the work for publication. We are grateful to Martine and Claude Boccara for their invaluable help with the interferometer device. We thank the INRA GeT-PlaGE platform (https://get.genotoul.fr/la-plateforme/get-plage) for sequencing the Epoisses virome and the INRA MIGALE bioinformatics platform (http://migale.jouy.inra.fr) for providing computational resources and support.

## SUPPLEMENTARY DATA

**Table S1**. Microbiological counts obtained for the nine cheese samples.

**Table S2**. DNA quantities obtained for the different samples studied.

**Table S3**. Comparison between phage titer and interferometry measure.

**Dataset S1**. Sequence comparison results of the 124 contigs composing the Epoisses virome.

**Dataset S2**. Predicted function and best BlastP hit for ORFs of Epvir4 and Epvir8.

**Fig. S1**. Functional annotation of the three most abundant viral contigs detected in the Epoisses cheese virome. Predicted functions of the ORFs identified in Epvir4 (A), 949_Epvir1 (B) and Epvir8 (C) are color-coded as follow: red for integrase, yellow for transcriptional regulation, orange for replication and recombination, green for DNA packaging and head, light blue for connector, dark blue for tail, light pink for homing endonuclease, dark pink for lysis, grey for hypothetical proteins, violet for additional functions. Some of the predicted functions of the ORFs are detailed for each contig.

**Fig. S2**. Comparison between phage concentration as measured by titration and using the interferometric light microscope (ILM). Six reference phages were used for the comparison, namely phages P1, T4, T5, T7, λCI857 and φX174.

**Fig. S3**. Epifluorescence image of a VLP solution obtained from Epoisses cheese. The VLP solution was diluted 10-fold in 1 ml of SM buffer with gluteraldehyde (2.5% final), incubated 10 minutes on ice and then flash frozen in liquid nitrogen. 100 μl of the nanoparticle solution was diluted into 5 ml of filtered water; and then filtered through a 25 mm anodisc membrane with pore sizes of 0.02 μm and stained with 100 μL of SYBR Gold (50×) for 15 min in the dark. Filters were placed on a microscopy slide with 100 μL of anti-fading solution Fluoromount, and observed immediately after preparation on a microscope Leica DMRA2 microscope equipped with a × 100 magnification oil-immersion objective and a COOLSNAP HQ camera (Roper Scientific, USA), using YFP filters. Images were captured and processed with METAMORPH V6.3r5.

